# Conditioned increase of locomotor activity induced by haloperidol

**DOI:** 10.1101/353912

**Authors:** Luis G. De la Casa, Lucía Carcel, Juan C. Ruiz-Salas, Lucía Vicente, Auxiliadora Mena

## Abstract

Dopamine antagonist drugs have profound effects on locomotor activity. In particular, the administration of the D2 antagonist haloperidol produces a state that is similar to catalepsy. In order to confirm whether the modulation of the dopaminergic activity produced by haloperidol can act as an unconditioned stimulus, we carried out two experiments in which the administration of haloperidol was repeatedly paired with the presence of distinctive contextual cues that served as a Conditioned Stimulus. Paradoxically, the results revealed a dose-dependent increase in locomotor activity following conditioning with dopamine antagonist (Experiments 1) that was susceptible of extinction when the conditioned stimulus was presented repeatedly by itself after conditioning (Experiment 2). These data are interpreted from an associative perspective, considering them as a result of a classical conditioning process.

## Introduction

An inherent characteristic of nature is change and, throughout the process of evolution, organisms endowed with a complex nervous system have developed psychological mechanisms that allow for anticipating these changes and producing responses that facilitate adaptation to the environment. One such mechanism is that of classical or Pavlovian conditioning, which has been proposed as a fundamental process to explain how organisms learn to respond adaptively in anticipation of the occurrence of environmental events (1,2). In fact, there are numerous examples that illustrate the relevance of classical conditioning in the field of the study of emotional processes (3,4), in the acquisition of eating habits (5,6) or its usefulness for the analysis and treatment of certain pathologies (7,8), among many others.

Another area in which the adaptive relevance of Pavlovian associations has been demonstrated is related to the effects of repeatedly presenting a neutral stimulus accompanied by the effects of a drug. This procedure has led to seemingly contradictory results, since while in some cases the Conditioned Response (CR) that appears has been similar to that produced by the drug (9,10), on other occasions the CR has been of an opposite nature to that induced by the drugs (11,12). Eikelboom & Stewart (1982) have proposed that the origin of these differences could be related to the effect of the drug on the nervous system: whilst on some occasions the Conditional Stimulus (CS) is associated with an Unconditioned Response (UR) dependent on the central nervous system, at other times the CS is associated with a peripheral UR that will appear to compensate for the central effects of the drug. In the first case, the association between the drug and the CS would lead to the appearance of a CR similar to the one that is produced by the drug, while in the second case, the CR would be opposite to that produced by the drug at the central level.

The first experimental evidence to highlight the effects of classical conditioning in the field of drugs was described by Pavlov himself, who reported that the repeated administration of morphine in the presence of a given context gave rise to a CR similar to that produced by morphine alone (14). From these pioneering studies, which demonstrated that contextual cues can be used as CSs that acquire the ability to induce physiological and behavioral states similar to those produced by the drug, a number of studies have been developed to demonstrate the conditioning of various responses produced by a wide range of drugs including, for example, morphine-induced hyperthermia (15,16), stereotypy or hyperactivity induced by amphetamine, cocaine, or apomorphine (17–20), amphetamine-induced hyperthermia (21), or haloperidol-induced catalepsy (22–24). The conditioning process supported by these drugs has been used to identify the neurobiological bases of learning (19), and has been considered as a possible relevant factor in the relapse of addicts (25,26), since it helps to explain the development of tolerance and the sensitization of drug-induced responses (27,28).

In our work, we will focus specifically on the conditioning of locomotor activity, using the administration of the dopaminergic antagonist haloperidol as a US. The usual procedure employed in this type of experiment involves a design that includes two groups that differ in terms of the time at which the drug is administered (29). For the first of the groups, which is usually referred to as the Paired group, the drug is administered before introducing the animal into the experimental context that will serve as the CS. After spending a period of time that usually ranges between 30 and 60 min in the context CS, an innocuous solution is administered and the animals are returned to their home cages. The second group, usually called the Unpaired group, first receives the saline solution in such a way that exposure to the context takes place in the absence of the drug, and the corresponding dose of the drug is administered before returning the animal to its home cage. After a rest period of around two days without receiving any type of drug or behavioral treatment, a test trial is carried out in which all animals of both groups are injected with the innocuous solution before introducing them to the context CS to record the activity.

Using this basic procedure, results have consistently revealed the existence of the conditioning of locomotor responses using dopamine agonists such as amphetamine or apomorphine. In particular, a significant increase in conditioned locomotor responses has been observed on the test trial for the paired group in comparison with the unpaired group (29–34). Less consistent are the results that have been obtained when the US employed is the dopamine antagonist haloperidol (35,36), possibly due to the fact that, depending on the dose administered, haloperidol can result in both an increase as well as a decrease in locomotor activity. More specifically, when a low dose of haloperidol (less than 0.1 mg / kg) is administered repeatedly, a progressive increase in the locomotor response is observed, which has been interpreted as the result of a sensitization process due to the selective blockade of the presynaptic autoreceptors that results in dopamine levels rising, leading to an increase in locomotor activity (36). However, when a higher dose is repeatedly administered (from 0.1 mg / Kg.), both pre and post-synaptic receptors are blocked, resulting in a reduction in locomotor behavior. This can even induce a state of catalepsy, in which the animals maintain unusual postures for prolonged periods of time (36–38). When, after repeated administration of the dopaminergic antagonist, a drug-free test is carried out in the same context in which the drug was administered, different results emerge depending on the dose of drug given during the conditioning trials. Thus, with doses of haloperidol that can be classified as high (specifically 0.1, 0.25 and 0.5 mg / Kg), an increase in conditioned catalepsy has been found in the Paired group with respect to the Unpaired control group on the test trial without the drug (38). However, Dias et al (2012) found an increase in locomotor activity in a group that had received ten pairings of the context-CS and a low dose of haloperidol (0.03 mg / kg), although in this case the subjects had received 5 trials in which 2.0 mg / kg of apomorphine had been injected before the conditioning test.

In the present study we set out to analyze the conditioning of locomotor activity following the repeated pairing of a context-CS and the effects of the administration of the dopaminergic D2 antagonist haloperidol. The method most commonly used in the literature to evaluate such behavioral patterns is either to observe movements in a limited space, generally an open field cage where the total distance traveled, the number of turns, grooming, etc. are usually recorded (39) or the so-called “bar test”, consisting in place the forepaws of the animal on a bar situated at a height adequate to the animal tested and record the time elapsed until the animal put down the paws on the floor (40). However, in our case we recorded the percentage of time that the animal remained in motion during each of the experimental sessions (60 min duration) since we expected a reduction in motor activity both after drug administration and when testing conditioning.

On the basis of previous findings, we anticipate that, with the concentrations of the drug we have used, after pairing the context with a dopaminergic antagonist (haloperidol) it will be observed a conditioned decrease in general activity (9,36–38).

## Experiment 1

The purpose of this experiment was to examine the conditioning of the locomotor response induced by the effect of two different doses of a drug that acts as a dopaminergic antagonist (haloperidol, 0.5 and 2.0 mg / kg). For this, the animals in the Paired condition received the administration of haloperidol before exposure to an experimental context that was to serve as a CS, whilst animals in the Unpaired condition received haloperidol after exposure to the experimental context.,

Based on the previous results we anticipate that with the selected doses (0.5 and 2.0 mg / Kg,) there will be a decrease in the activity on a subsequent drug-free test trial in presence of the conditioned context that probably will be more intense with the higher dose.

### Subjects

32 male Wistar rats (n=8) experimentally naïve, participated in this experiment. Mean weight at the start of the experiment was 384 g. (range 292 - 490). Food and water were available <u>ad libitum</u> throughout the experiment. Animals were individually housed and maintained on a 12:12 h light:dark cycle (lights on at 06:00 h). All behavioral testing was conducted during the light period of the cycle. Four days before the start of the experimental sessions, each of the animals was handled 5 min daily. All procedures were conducted in accordance with the guidelines established by the European Union Council established by the Directive 2010/63/EU, and following the Spanish regulations (R.D 53/02013) for the use of laboratory animals. The ethical commission of University of Seville supervised and approved all the procedures and all protocols used in the this specific study (report: CEEA-US2015-28/4)

### Apparatus and Materials

Four identical Panlab conditioning boxes (model LE111, Panlab/Harvard Apparatus, Spain) were used, each measuring 26 × 25 × 25 cm (H × L × W). Each chamber was enclosed in a sound-attenuating cubicle (model LE116. Panlab/Harvard Apparatus, Spain). The walls of the experimental chambers were made of white acrylic. A loudspeaker located at the top of each chamber produced a 70 dB 2.8-kHz noise used as background, and the floor consisted of stainless steel rods, 2 mm in diameter, spaced 10 mm apart (center to center). Each chamber rested on a platform that recorded the signal generated by the animal movement through a high sensitivity Weight Transducer system. Such signal was automatically converted into percent of general activity, defined as the percentage of the total time that movement was detected on 2-min periods, by a commercial software (StartFear system software, Panlab/Harvard Apparatus, Spain). Sampling was performed continuously at a frequency of 50Hz.

Haloperidol (Pensa Pharma) dissolved in 0.1% ascorbate/saline (2.0 mg/ml) was injected subcutaneously in the nape of the neck at a dose of 0.5 or 2.0 mg/kg. A 0.1% ascorbate/saline solution was used as vehicle. A delay of 20 min was introduced from the drug administration to the introduction of the animals in the experimental chambers.

### Procedure

Four groups were arranged following a 2 × 2 factorial design, with main factors Conditioning (Paired vs. Unpaired) and Dose (0.5 vs. 2.0 mg/Kg of haloperidol). Regarding the Conditioning factor, those animals in the Paired condition received an injection of the correspondent drug before to be introduced in the experimental context, and an injection of vehicle before to be returned to their home cages; those rats in the Unpaired condition received the vehicle before experimental context exposure, and the drug after each session (and before to be returned to the home cages).

The experimental treatment started with a single 60-min. baseline session intended to measure general activity of each animal without the effect of the drug, and to habituate the rats to the new context (before this session each animal was injected with vehicle). The next day started the context conditioning stage. This phase comprised four 60-min sessions conducted on consecutive days. Those animals in the Paired/0.5, and Paired/2.0 groups were injected with the correspondent haloperidol dose before being introduced on the experimental context. Immediately after each session, each animal was injected with an equivalent dose of vehicle before to return to the home cage; those animals in the Unpaired/0.5, and Unpaired/2.0 groups received the Vehicle before context exposure, and the drug just before to be returned to their home cages. Mean percent of activity was registered for each conditioning session as an index of sensitization.

A test session was conducted 48 hours after the last conditioning day, and consisted in injecting the corresponding dose of vehicle for all rats and registering activity for 60 min. in presence of the experimental context in periods of 10 min. The dependent variable used as an index of conditioning was mean percent of activity.

## Results

### Baseline

Mean percent activity during the baseline day collapsed across 60 min was 52.24% (range 28.99 % - 73.94 %). A 2 × 2 ANOVA (Conditioning × Dose) conducted on mean activity revealed that neither the main effects nor the interaction was significant (all *p*s>.40).

### Context conditioning

Fig. 1 shows mean activity across the four conditioning days as a function of groups. As can be seen in the figure, those animals that were injected with haloperidol before context exposure (Paired condition) showed a very low and stable percent of activity during all conditioning days. Those animals injected with the drug after context exposure (Unpaired condition) showed higher levels of activity that decreased across days, probably reflecting a habituation process.

**Fig. 1.**
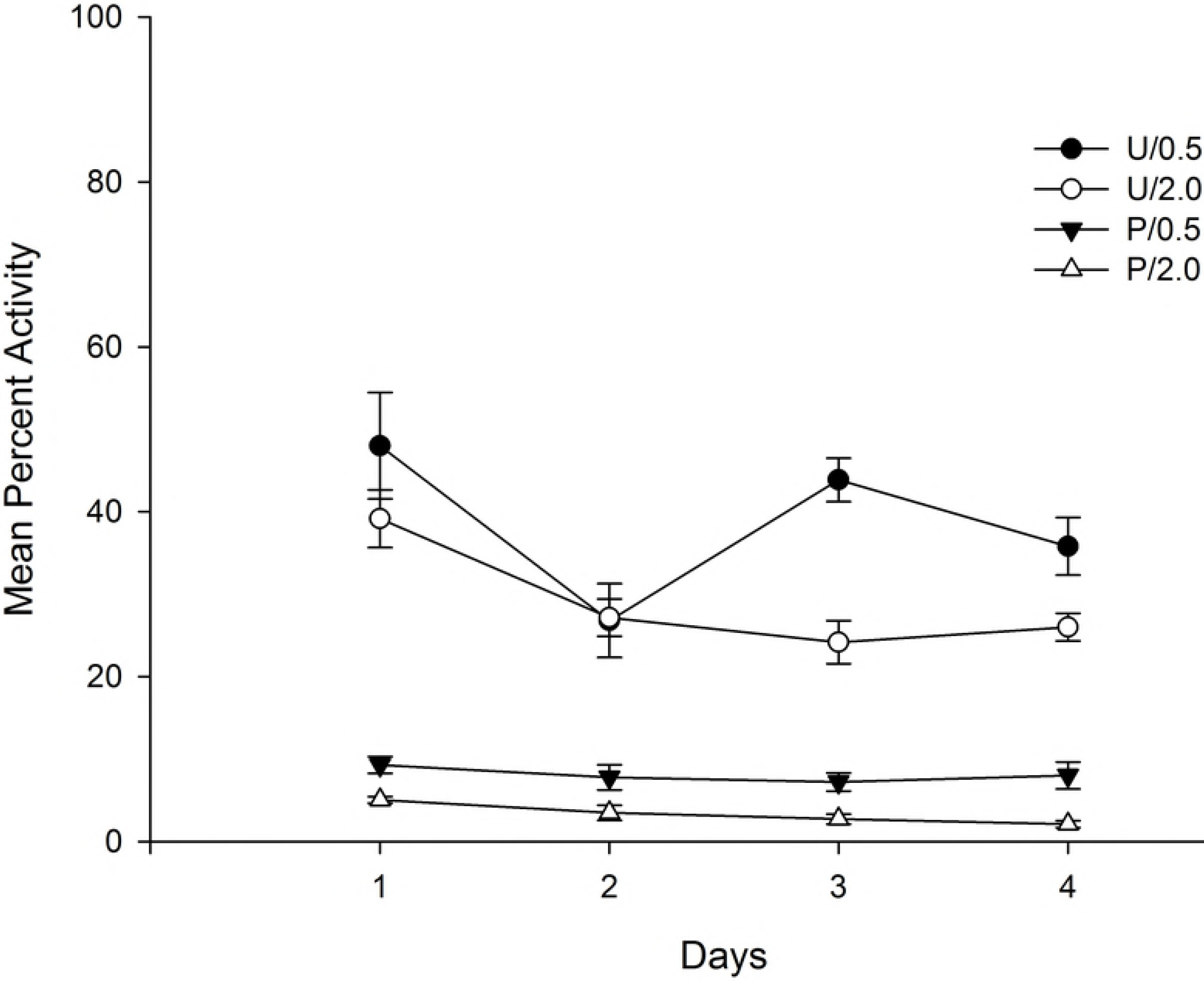
Mean percent activity on conditioning days as a function of conditioning, and haloperidol dose. Percent activity was collapsed across each 60 min session. The animals had received either 0.5 mg/Kg or 2.0 mg/Kg of haloperidol before (P: Paired) or after (U: Unpaired) being introduced in the context-CS for 60 min.

A 4 × 2 × 2 mixed ANOVA with main factors Days (within-subject), Conditioning (Paired vs. Unpaired), and Dose (0.5 vs. 2.0) was conducted on mean percent activity collapsed across each 60 min session. The main effects of Days and Conditioning were significant, *F*(3,84)=9.21; *p*<.001; η^2^=25, and *F*(1,28)=357.90; *p*<.001; η^2^=93, respectively. The main effect of Days reflects a general reduction of activity across sessions, and the effect of Conditioning was due to the overall lower levels of activity for those animals in the Paired as compared to those in the Unpaired condition (Mean = 5.02%, SD = 3.35, and Mean = 35.74%, SD = 8.90, respectively). The main effect of Dose was also significant, *F*(1,28) = 22.86; *p*<.001; η^2^=45, due to a higher level of activity for those animals that received the 0.5 mg/kg as compared to those injected with the 2.0 mg/kg (Mean = 24.54%, SD = 19.31, and Mean = 16.22%, SD = 13.56, respectively). Finally, the Days x Conditioning interaction was significant, *F*(3,84) = 5.95; *p*<.01; η^2^=.18, reflecting a progressive reduction of activity across days that was restricted to those animals that received the vehicle injection before context exposure. No more interactions were significant (all ps>.06).

### Test

Fig. 2 (panel A) shows mean percent activity during the test day collapsed across 10-min periods as a function of Conditioning (Paired vs. Unpaired) and haloperidol Dose (0.5 vs. 2.0 mg/Kg). Fig. 2 (panel B) depicts mean activity collapsed across the entire session duration as a function of Conditioning and Dose. As can be seen in the upper panel of the figure, mean percent activity decreased across the test session, but it was higher for the animals in the Paired/0.5 Group. Similarly, as can be seen in the bottom section of Fig. 2, there was a general increase in activity that was restricted to the group that had received the lower dose of the drug before context exposure (Paired/0.5) as compared to the group that had received the drug after context exposure (Unpaired/0.5).

**Fig. 2.**
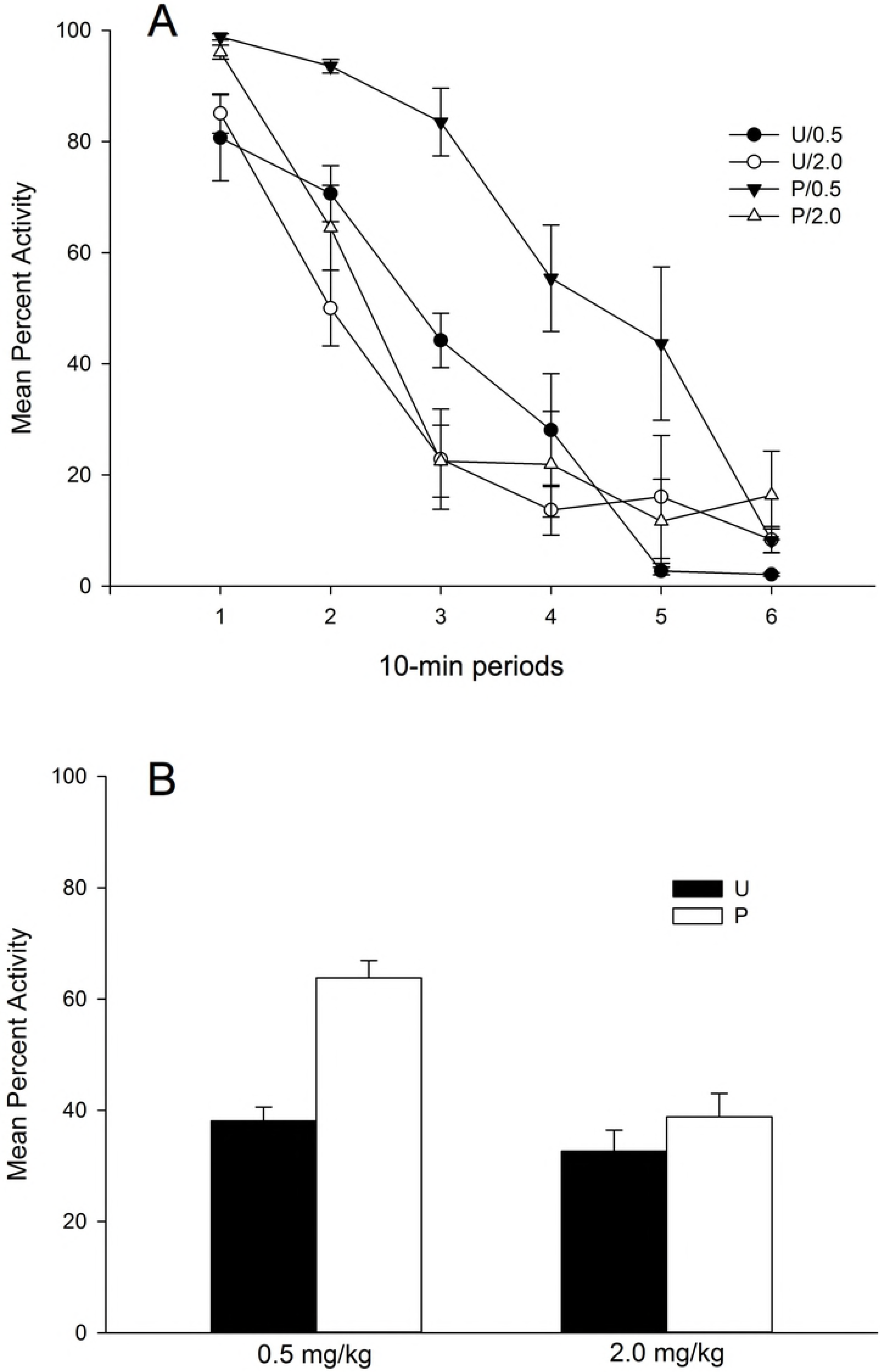
Mean percent activity on the drug-free test day as a function of Conditioning, and haloperidol Dose. (A) Percent activity is represented collapsed across 10-min periods, and (B) across the complete 60-min session. The animals had received 0.5 mg/Kg of haloperidol during the four days of the conditioning stage before (condition P: Paired) or after (condition U: Unpaired) been exposed for 60 min to the context-CS. Test session was drug-free.

A 6 × 2 × 2 ANOVA with main factors 10-min Periods (within-subjects), Drug, and Conditioning performed on mean percent activity collapsed across 10 min periods revealed a significant main effect of Periods, *F*(5,140)=91.34; *p*<.001; η^2^=77, due to the overall reduction of activity across the session, and a significant Periods × Drug interaction, *F*(5,140)=7.66; p<.001; η^2^=21, that reflects a faster decrease of activity across 10-min periods for the animals that received 2.0 mg/kg as compared to those that received the 0.5 mg/kg haloperidol dose. No more interactions involving the Periods factor were significant (ps>.06). The analyses involving the between-subject factors revealed significant main effects of Dose and Conditioning, *F*(1,28)=19.47; *p*<.001; η^2^=.41, and *F*(1,28)=21.47; p<.001, η^2^=43, respectively. The main effect of Dose reflects a significant higher percent of activity for those animals injected with the 0.5 mg/Kg dose as compared to those injected with the 2.0 mg/Kg. dose (mean = 50.94%, SD = 15.38, and mean = 35.73%, SD = 11.33, respectively). The main effect of Conditioning reflects an overall higher level of activity for the rats in the Paired as compared to those in the Unpaired condition (mean = 51.33%, SD = 16.36, and mean = 35.35%, SD = 9.2, respectively).

Importantly, the Conditioning × Dose interaction was significant, *F*(1,28)=8.09; *p*<.01, η^2^=.22. <u>Post-hoc</u> comparisons comparing groups (Bonferroni, *p*<.05) revealed that the association between the context and the effect of the drug resulted in an increased activity at testing as indicated for a significant difference between Paired/0.5 vs. Unpaired/0.5 groups, However, the effect of context conditioning did not appear when the 2.0 mg/Kg dose was injected, since there were no significant differences between Paired/2.0 vs. Unpaired/2.0 groups. Also, percent of activity was higher in the Paired/0.5 as compared to the Paired/2.0 group, but there were no differences between groups in the Unpaired condition.

## Experiment 2

The results of Experiment 1 revealed that after four pairings of a 0.5 mg/kg dose of haloperidol with an initially neutral context, the latter acquired the ability to induce an increase in the overall activity of the animals on a drug-free test trial. A possible explanation for this result from a non-associative perspective is that haloperidol in the Paired condition had impeded proper processing of the context during conditioning stage. Therefore, the context would have been functionally novel at time of testing and it would have elicited non-habituated exploration responses. However, such interpretation can be ruled out since the same result should have appeared in the animals injected with the higher dose of haloperidol before context exposure, but locomotor activity was similar at testing when comparing Paired/0.2 vs. Unpaired/0.2 groups.

Since this result not only fails to support our initial hypothesis, but also goes in the opposite direction, we designed an additional experiment to replicate it, and to test if a manipulation that typically affects to the CR affects to the predicted increase in locomotor activity (an extinction procedure). Therefore, in the following experiment, two groups were used that received exactly the same treatment described for the Paired/0.5 and Unpaired/0.5 groups in Experiment 1, but four free-drug test trials were programmed in order to evaluate the effect of an extinction process on the CR. Considering the results of the first experiment, we now expect to find a conditioned increase in locomotor activity in the test phase for the Paired group when compared with the Unpaired group, and a decrease of such response across extinction days.

### Subjects

16 male Wistar rats (n=8) experimentally naïve, participated in this experiment. Mean weight at the start of the experiment was 339 g. (range 459 - 266). The animals were housed and maintained as described for Experiment 1.

### Apparatus, materials and procedure

The apparatus, materials, and procedure were the same described for the groups Paired/0.5 mg/kg, and Unpaired/0.5 mg/kg in Experiment 1, except that four free-drug tests trials, instead of one, were conducted after conditioning stage.

## Results

### Baseline

Mean percent of activity on the baseline day was 47.37 % (Range: 25.82% - 65.67%). A one-way ANOVA conducted on mean percent activity as a function of Groups revealed that the differences were non-significant (*F*<1).

### Context conditioning

Fig. 3 depicts mean percent of motor activity collapsed across the 60 min for the four conditioning days as a function of Groups (Paired vs. Unpaired). As can be seen in the figure, the rate of activity was low and constant across the conditioning days for the Paired Group. The animals in the Unpaired Group showed a high percentage of motor activity that decreased across conditioning days reflecting the habituation to the contextual cues.

**Fig. 3.**
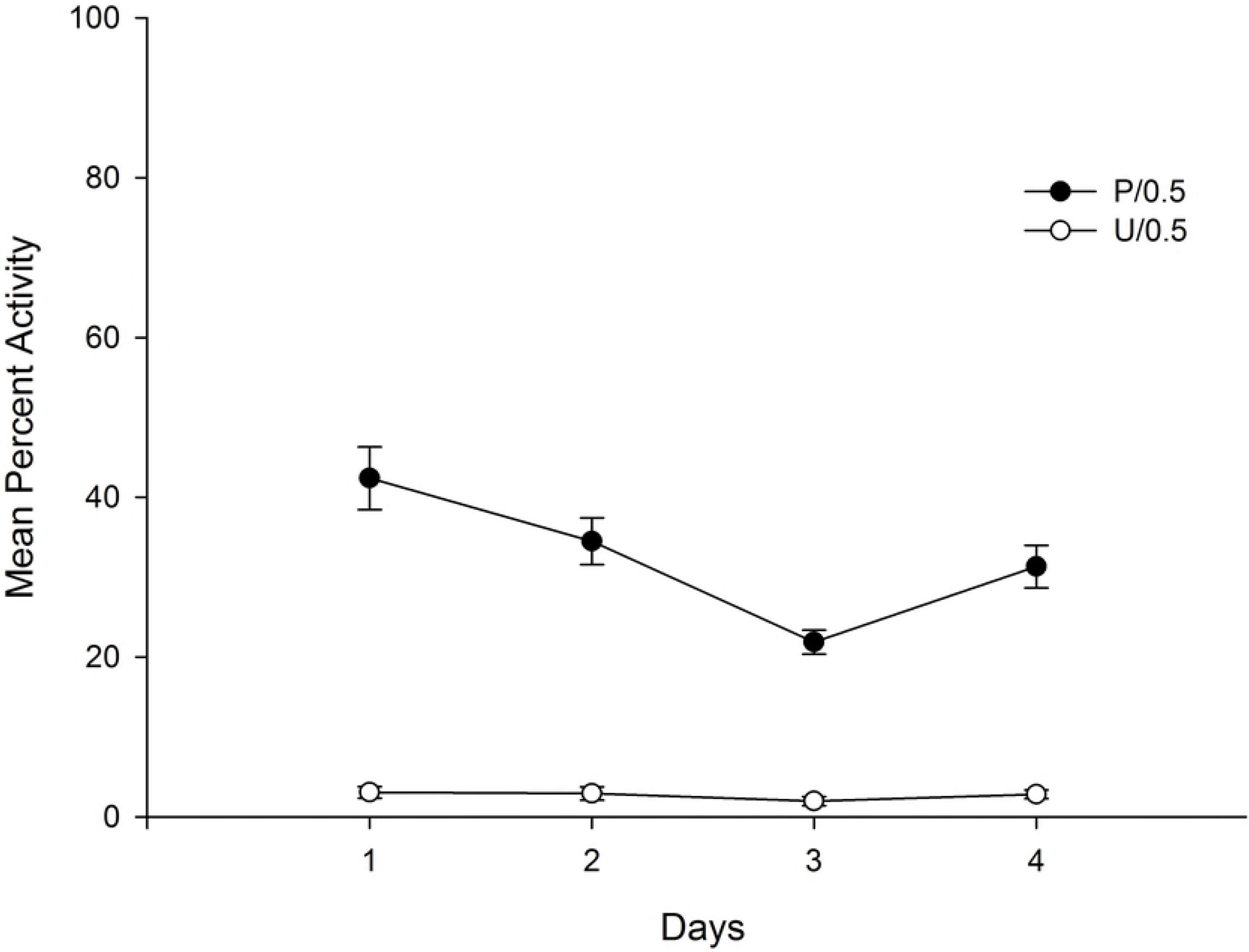
Mean percent activity on conditioning days as a function of conditioning. Percent activity was collapsed across each 60 min session. The animals had received 0.5 mg/Kg of haloperidol before (P: Paired) or after (U: Unpaired) being introduced in the context-CS for 60 min.

A 4 × 2 mixed ANOVA with main factors Days (within-subject) and Group (Paired vs. Unpaired) was conducted on mean percent activity for each day (collapsed across the 60 min of each trial duration). The main effect of Group was highly significant, *F*(1,14)=288,64; *p*<.001; η^2^=95, due to the decrease in activity for those animals in the Paired as compared to those in the Unpaired Group. This result confirmed the effectiveness of haloperidol to reduce locomotor activity. The main effect of Days, and the Days x Groups interaction were also significant, *F*(3,42) = 13.20; *p*<.001; η^2^=49, and *F*(3,42) = 10.65; *p*<.001; η^2^=43. The main effect of Days reflects the overall decrease of activity across days, and the interaction was due to a progressive reduction of activity across days for the animals in the Unpaired Group (due to habituation) that contrast with the low and constant activity for the rats in the Paired Group.

### Test

The top section of Fig. 4 shows mean percent activity during the first test day collapsed across 10-min periods as a function of Conditioning (Paired vs. Unpaired), and the bottom section of the figure depicts mean percent of activity collapsed across the four 60 min free-drug extinction days for the Paired and the Unpaired Groups. An inspection of the upper section of the figure reveals that motor activity remained higher during all the 10-min periods in the Paired as compared to the Unpaired Group. In addition, and as can be seen in the lower section of Fig. 4, there was a progressive decrease of locomotor activity across days in the Paired Group that can be interpreted as a result of the extinction process.

**Fig. 4.**
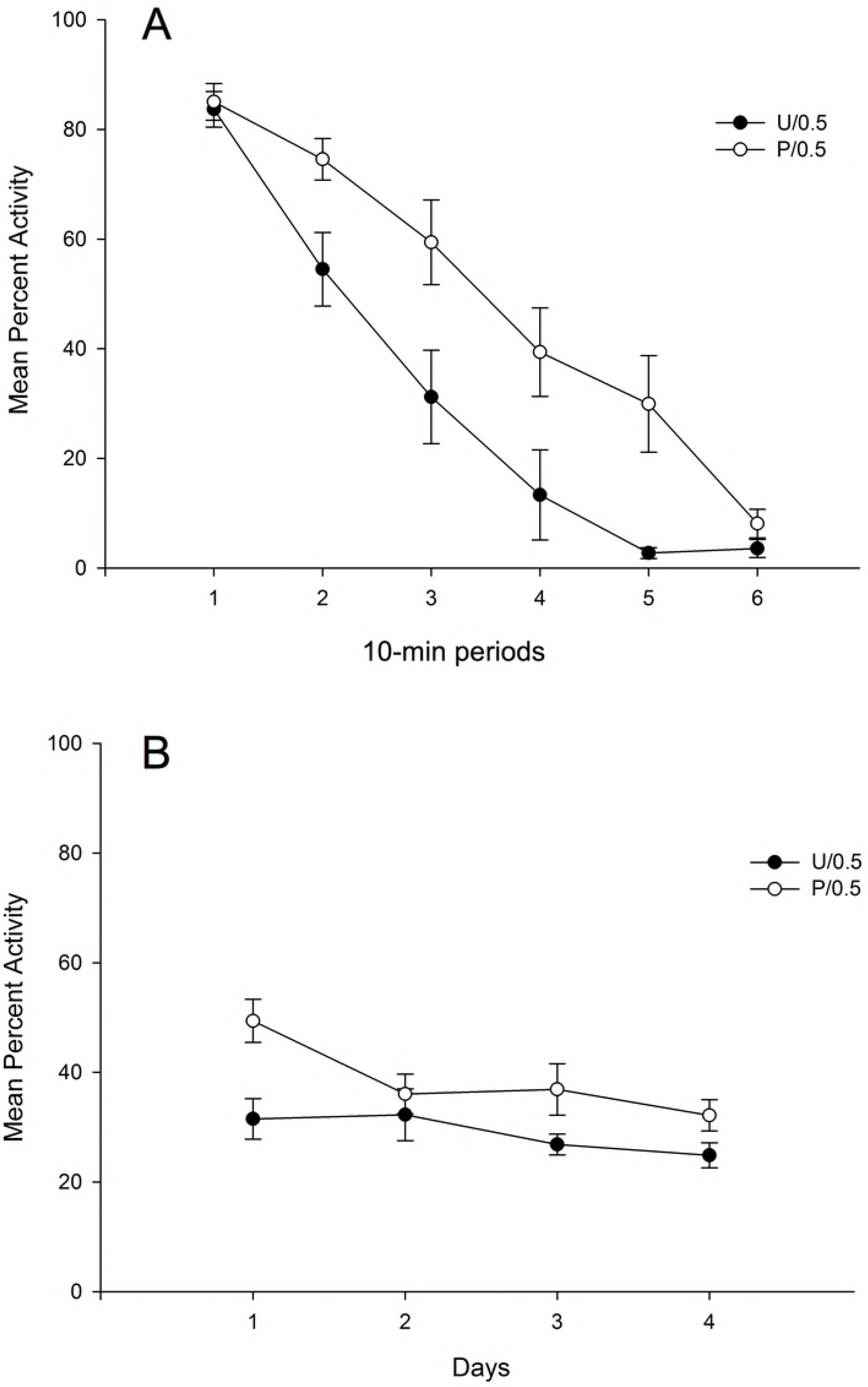
Mean percent activity on the drug-free test day as a function of Conditioning, and haloperidol Dose. (A) Percent activity is represented collapsed across 10-min periods, and (B) across the complete 60-min session. The animals had received 0.5 mg/Kg of haloperidol during the four days of the conditioning stage before (condition P: Paired) or after (condition U: Unpaired) been exposed for 60 min to the context-CS. Test session was drug-free.

A 6 × 2 mixed ANOVA with main factors Periods (within-subjects) and Group performed on mean percent activity collapsed across 10 min periods for the first free-drug test day revealed a significant main effect of Period, *F*(5,70)=69.87; *p*<.001; η^2^=.83, due to the overall reduction of activity across the session, and a significant Period × Group interaction, *F*(5,70)=2.77; *p*<.05; η^2^=17, due to a faster decrease of activity across 10-min periods for the animals in the Unpaired as compared to those in the Paired Group. The main effect of Group was also significant, *F*(1,14)=10.95; *p*<.01; η^2^=.44, reflecting the higher level of activity for the rats in the Paired as compared to those in the Unpaired condition (mean = 49.39%, SD = 10.46, and mean = 31.49%, SD = 11.17, respectively). This result replicates the conditioned increase of locomotor activity obtained in the Paired/0.5 Group from Experiment 1.

Additionally, a 4 × 2 mixed ANOVA with main factors Days and Group was performed on mean percent activity in order to test whether the extinction procedure was effective in reducing the CR. The analysis revealed a significant main effect of Group, *F*(1,14)=5.24; *p*<.05; η^2^=27, due to the overall conditioned increase in activity showed for those animals in the Paired as compared to the Unpaired Group. The main effect of Days was also significant, *F*(3,42)=9.82; *p*<.001; η^2^=41, due to a general decrease of locomotor activity across days. Finally, the 2-way interaction was significant, *F*(3,42)=3.50; *p*<.05; η^2^=20, due to the progressive decline in locomotor activity for the Paired Group reflecting the extinction of locomotor conditioning across days.

## General discussion

The results of Experiment 1, in which two different doses of the dopaminergic antagonist haloperidol were administered, were not consistent with the hypothesis of the conditioning of drug-induced locomotor activity from which we anticipated a decrease in activity in the presence of the CS that had been paired with the dopaminergic antagonist (36,37,41). Firstly, none of the doses given to the Paired groups produced an effect of sensitization to the drug, since for these animals the percentage of activity remained at low and constant levels from the first day. It is possible that our dependent variable (the general activity of the animal) was not sufficiently sensitive to repeated administrations of the drug, since in the experiments in which this sensitization effect has been observed, other indices of activity have been used. In contrast, in the test phase a significant increase in locomotor activity was observed for those animals in the Paired condition that had received the lowest dose of haloperidol (0.5 mg / Kg) with respect to the Unpaired control group. This same result was replicated in Experiment 2 that also revealed that the conditioned increase of activity was affected by an extinction treatment.

In view of these results, we can conclude that the repeated pairing of a neutral stimulus (in our case the experimental context) with the administration of a 0.5 mg / Kg dose of haloperidol produces a conditioned increase in locomotor activity during a subsequent test phase conducted in the experimental context. Some authors have proposed that this type of response could be the result of a non-associative process, since the administration of dopaminergic agonist drugs prior to exposure to the context could hinder the processing of the latter, so that the context would be functionally new at the time of testing and would thus elicit the same orientation responses that would be expected in response to a novel context (42,43). However, this possibility can be ruled out attending to the results of the groups that received a 2.0 mg/Kg dose of haloperidol in Experiment 1, since from this perspective the higher dose of haloperidol should have induced a similar o even a bigger increase in locomotor activity at testing as that observed in the Group that received the 0.5 mg/Kg. dose. However, there were no significant differences between the Paired vs. the Unpaired Group that received the highest antagonist’s dose, indicating that the hypothetical reduced processing of the context during the conditioning stage can not explain the increased activity observed during testing.

A second possibility that has been proposed to explain the conditioning of locomotor activity is related to the rewarding properties of dopaminergic agonist drugs, which, after being paired with the context, would allow the latter to evoke approach responses that would be manifest during the conditioning test as an increase in locomotor activity (44,45). This account, which links the association between the context and the effects of the drug with a reward-related incentive learning process, takes into account the rewarding value of the drugs that have usually been used in these types of experiments (such as amphetamine, apomorphine, and cocaine), which is a consequence of an increase in dopaminergic activity in the mesotelencephalic reward system (46). This hypothesis, however, could not explain our results, since the drug administered was a dopaminergic antagonist that has no rewarding action (47) and that has even proven to be effective in blocking the reinforcing value of certain stimuli or drugs with hedonic value (48,49).

A third account of the origin of the increase in locomotor activity observed after pairing the context with a drug can be established in strictly Pavlovian terms, based on the assumption that the CS is a stimulus that acquires the same properties as the US and, therefore evokes the same type of responses after the conditioning process (50,51, but see 52). This theory of stimulus substitution (14) seems to be at a first glance difficult to conciliate with our results, since the observed CR (an increase in locomotor activity) is opposite to the UR (a reduction of locomotor activity). However, the fact that the repeated administration of a low dose of haloperidol has proved effective in inducing an increase in locomotor activity (36) makes it possible to reconcile our results with this classical conditioning perspective.

More specifically, there is ample evidence to suggest that the main pharmacological action of haloperidol consists of the blockade of D2 receptors, some of which are autoreceptors located in terminals and dopaminergic dendrites, while others are located postsynaptically in the soma, dendrites, and terminals of noradrenergic neurons (53). Haloperidol at medium or high doses, by blocking presynaptic D2 receptors (autoreceptors), increases the release of dopamine (54), but the increase in dopaminergic transmission is nullified by the blockade of post-synaptic D2 receptors. However, at low doses, haloperidol exerts its antagonist action only in the autoreceptors, and not by blocking the post-synaptic receptors, since the concentration of drug required to produce antagonist action in the post-synaptic site would be greater (55). As we have indicated above, Dias et al (2012) showed that the repeated administration of a very low dose of haloperidol (0.03 mg / Kg) produces an increase in locomotor activity that could be related to the selective blockade of presynaptic autoreceptors. In the same study, a high dose of haloperidol (1.0 mg / Kg) caused an inhibitory effect on locomotion that could be related to the blockade of post-synaptic D2 receptors.

Albeit speculative, based on the fact that in our experiments with haloperidol we used only 4 pairings of the context-CS and the drug-US, compared to the 8 pairings in the experiments of Schmidt & Beninger (2006), or the 10 employed by Banasikowsky & Beninger (2012) or Dias et al. (2012), and taking into account that the CR is usually of a lower intensity than the UR (56,57), we suggest that in our experiments the presentation of the context associated with the 0.5 mg / kg dose of haloperidol may have led to a low intensity CR that would have been functionally equivalent to the response induced by a low dose of haloperidol. This CR could have blocked the presynaptic dopamine autoreceptors, preventing the feedback mechanism that would limit the release of the neurotransmitter, while the postsynaptic receptors would not have been affected, leading to an excessive dopamine reuptake that could have caused the conditioned increase in locomotor activity.

## Acknowledgments

The authors thank to Félix Hermoso and Francisco José Pérez Díaz for their useful comments and help in running the experimental sessions.

